# Transcranial ultrasound stimulation in humans is associated with an auditory confound that can be effectively masked

**DOI:** 10.1101/2020.03.07.982033

**Authors:** Verena Braun, Joseph Blackmore, Robin O. Cleveland, Christopher R. Butler

## Abstract

**Background:** Transcranial ultrasound stimulation (TUS) is emerging as a potentially powerful, non-invasive technique for focal brain stimulation. Recent animal work suggests, however, that TUS effects may be confounded by indirect stimulation of early auditory pathways.

**Objective:** We aimed to investigate in human participants whether TUS elicits audible sounds and if these can be masked by an audio signal.

**Methods:** In 18 healthy participants, T1-weighted magnetic resonance brain imaging was acquired for 3D ultrasound simulations to determine optimal transducer placements and source amplitudes. Thermal simulations ensured that temperature rises were <0.5 °C at the target and <3 °C in the skull. To test for non-specific auditory activation, TUS (500 kHz, 300 ms burst, modulated at 1 kHz with 50% duty cycle) was applied to primary visual cortex and participants were asked to distinguish stimulation from non-stimulation trials. EEG was recorded throughout the task. Furthermore, ex-vivo skull experiments tested for the presence of skull vibrations during TUS.

**Results:** We found that participants can hear sound during TUS and can distinguish between stimulation and non-stimulation trials. This was corroborated by EEG recordings indicating auditory activation associated with TUS. Delivering an audio waveform to participants through earphones while TUS was applied reduced detection rates to chance level and abolished the TUS-induced auditory EEG signal. Ex vivo skull experiments demonstrated that sound is conducted through the skull at the pulse repetition frequency of the ultrasound.

**Conclusion:** Future studies using TUS in humans need to take this auditory confound into account and mask stimulation appropriately.

## Introduction

Transcranial ultrasound stimulation (TUS) uses low intensity focused ultrasound delivered through the skull to cause direct modulation of neuronal function [1-4]. In animal studies, TUS has been shown to modulate activity in several brain areas, including sensorimotor regions, visual cortex, frontal eye fields, anterior cingulate cortex and thalamic targets, resulting in behavioural as well as electrophysiological changes [5-20]. Furthermore, longer term connectivity changes have been identified in non-human primates following offline TUS [4,21]. Several studies have shown that TUS can be applied safely to healthy human participants [22] to modulate behaviour and neural activity in brain regions including somatosensory, visual, and motor cortex as well as to deeper thalamic nuclei [23-28]. These data have resulted in TUS emerging as a safe, potent, non-invasive brain stimulation tool [1], with better spatial accuracy and greater depth than established techniques such as transcranial magnetic or electrical stimulation [29].

However, recent reports from rodent studies have suggested that behavioural and neural effects of TUS may in fact result from indirect widespread auditory activation [30,31]. In guinea pigs, strong responses in the primary auditory cortex were observed independent of the sonicated brain target and transection of the auditory nerve or removal of the cochlear fluid abolished the response [30]. In transgenic mice, activity produced by ultrasound bursts strongly resembled activity associated with audible sound and was eliminated by chemical deafening [31]. It has been shown in mice that auditory activation arises from sharp edges in a TUS rectangular envelope stimulus [32] and that the auditory activation could be eliminated by smoothing the onset and offset of a continuous wave stimulation over 12 ms. However, such long smoothing cannot be employed for the 0.5 ms pulses that are commonly used in TUS.

In neurostimulation research it is well accepted that confounding factors have to be carefully controlled [33] in order to ensure that the effects observed are indeed the result of having stimulated a certain brain area, rather than by extraneous effects such as somatosensory stimulation [34,35]. In TUS this is particularly important since the precise mechanisms by which neuromodulation occurs are not well understood, although these remain the subject of intense research efforts [36]. It is important therefore to determine whether TUS may also have auditory side-effects in humans that could impact outcomes.

Using stimulation parameters similar to those employed in previous human studies, we applied TUS for 300ms at 500kHz modulated with a 1kHz square wave (50 % burst duty cycle (BDC)) to the right visual cortex of 18 healthy human participants. Stimulation and sham trials were presented in randomised order and subjects were asked to distinguish trials with active stimulation from trials in which no stimulation was applied. EEG was recorded throughout. Participants were reliably able to hear sound during stimulation trials, and therefore we further investigated the nature of this phenomenon with ex vivo skull recordings and tested the behavioural and electrophysiological effectiveness of auditory stimulus masking.

## Material and Methods

### TUS-EEG

#### Participants

Participants were screened for contraindications against brain stimulation [37] and magnetic resonance imaging and did not report hearing impairments. They were part of a larger study looking into the effects of TUS on visual processing and were remunerated for their time. This manuscript reports data from 18 healthy human participants who completed this part of the study (11 female and 7 male, mean age 26.22 +/-7.25 years). Informed consent was acquired from every participant prior to the experiment. The study was approved by the University of Oxford Medical Sciences Interdivisional Research Ethics Committee.

#### Experimental Setup and Procedure

The experiment was split into two blocks (Fig. 1a,b) which were conducted in the same order by every participant. First, participants took part in the unmasked block. This part consisted of 100 trials (50 stimulation, 50 sham stimulation trials) which were presented in randomised order. Participants were asked to keep their eyes fixed at the centre of a computer screen. In stimulation trials, TUS was applied 2.7-3 s after fixation onset for 300 ms before a question mark appeared prompting participants to indicate whether they thought they were stimulated or not, using their right hand. This block was followed by the masked block in which a masking tone was played through earphones in every trial, approximately 112 ms prior to the onset of the ultrasound stimulation for 700 ms, before the question mark appeared prompting participants to indicate whether they thought they were stimulated or not. Similar to the unmasked condition, 100 trials split into 50 stimulation and 50 no stimulation sham trials were presented in randomised order. At the end of each experimental condition block, participants were asked whether they had experienced any positive visual phenomena (such as phosphenes).

**Fig. 1.**
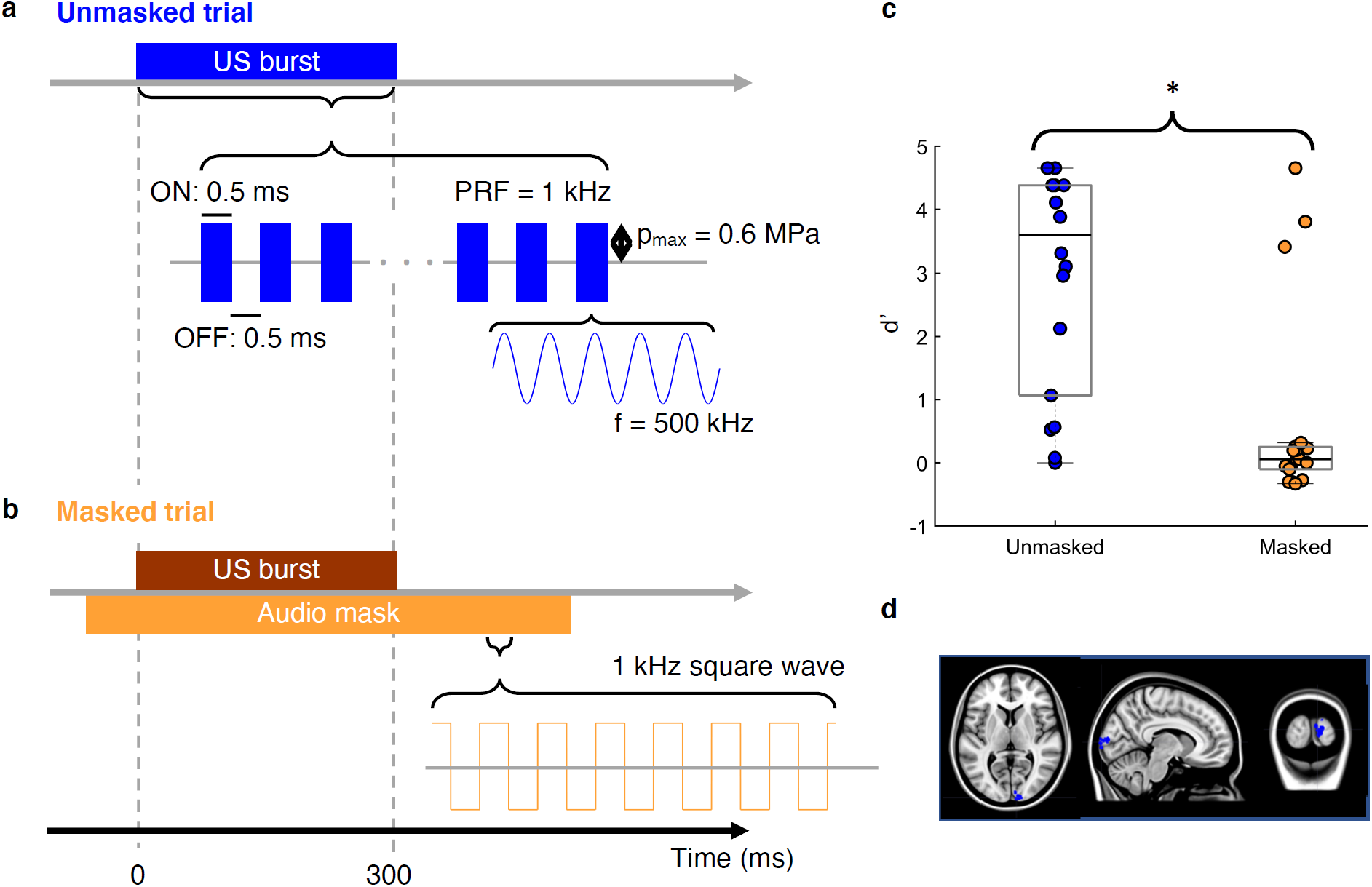
Experimental design and results. Participants performed a detection task in which they were asked to indicate whether a given trial was a stimulation or sham trial. The experiment was split into two conditions and was always administered in the same order starting with unmasked trials. **a, Unmasked trials**. In the unmasked condition, participants received TUS in half of the trials, while in the other half no TUS was applied. Each TUS trial consisted of a 300ms burst, which was made up of 0.5 ms pulses of 500 kHz ultrasound (250 cycles) alternating with 0.5 ms of no ultrasound (that is, a burst duty cycle (BDC) of 50 % and PRF of 1 kHz). The maximum modelled peak positive pressure at the target cortical site (V1) was 0.6MPa. **b, Masked trials**. The masked condition comprised the same TUS stimulation protocol but with the addition of an audio mask delivered via earphones to the participants. The audio mask was applied during TUS and sham trials in the masked condition only. The audio mask consisted of a 1 kHz square wave starting approximately 110ms before TUS onset and lasting 700 ms. **c, Behavioural results**. Boxplots depicting d’ values for unmasked and masked trials. Black lines indicate the median d’ value and filled circles correspond to individual data points. Detection rates were significantly lower in the masked condition compared to the unmasked condition. Three participants continued to detect stimulation in the presence of the masking signal. **d, Stimulation sites**. Stimulation sites shown in MNI space. Target brain sites were identified visually. Sites were chosen in the right visual cortex that could be reached easily with TUS.

Prior to the main experiment, T1-weighted high-resolution images were acquired from every participant with a Siemens 3T Trio system. For TUS modelling, tissue types were segmented using MARS [38] for SPM 8 (Wellcome Department of Cognitive Neurology, London. UK; www.fil.ion.ucl.ac.uk/spm) and target brain sites were identified visually in the right visual cortex (Fig. 1d).

Both acoustic and thermal participant-specific modelling was carried out to determine appropriate source locations and amplitudes in order to focus to the intended targets as well as satisfy the safety constraints (maximum peak modelled pressure in CSF or brain tissue = 0.6 MPa, maximum temperature rise in skull bone = 3 °C, maximum temperature rise in brain tissue = 1 °C) similar to other studies involving human subjects (see Fig 2 of Ref [36]). Numerical modelling was carried out using k-Wave, a pseudospectral time domain solver [39]. For the acoustic simulations, the skull bone was treated as a homogeneous medium and an effective fluid with transducer positions and orientations selected based on the phase distributions obtained from back-propagation of an US source positioned at the target to the source. Subsequent forwards simulations were then conducted to determine the in situ pressure fields and amplitudes, and hence determine appropriate source intensities. Finally, thermal simulations were carried out to calculate the associated temperature rises based on the estimated pressure fields. More details on the modelling protocol can be found in Blackmore et al. [40].

**Fig. 2.**
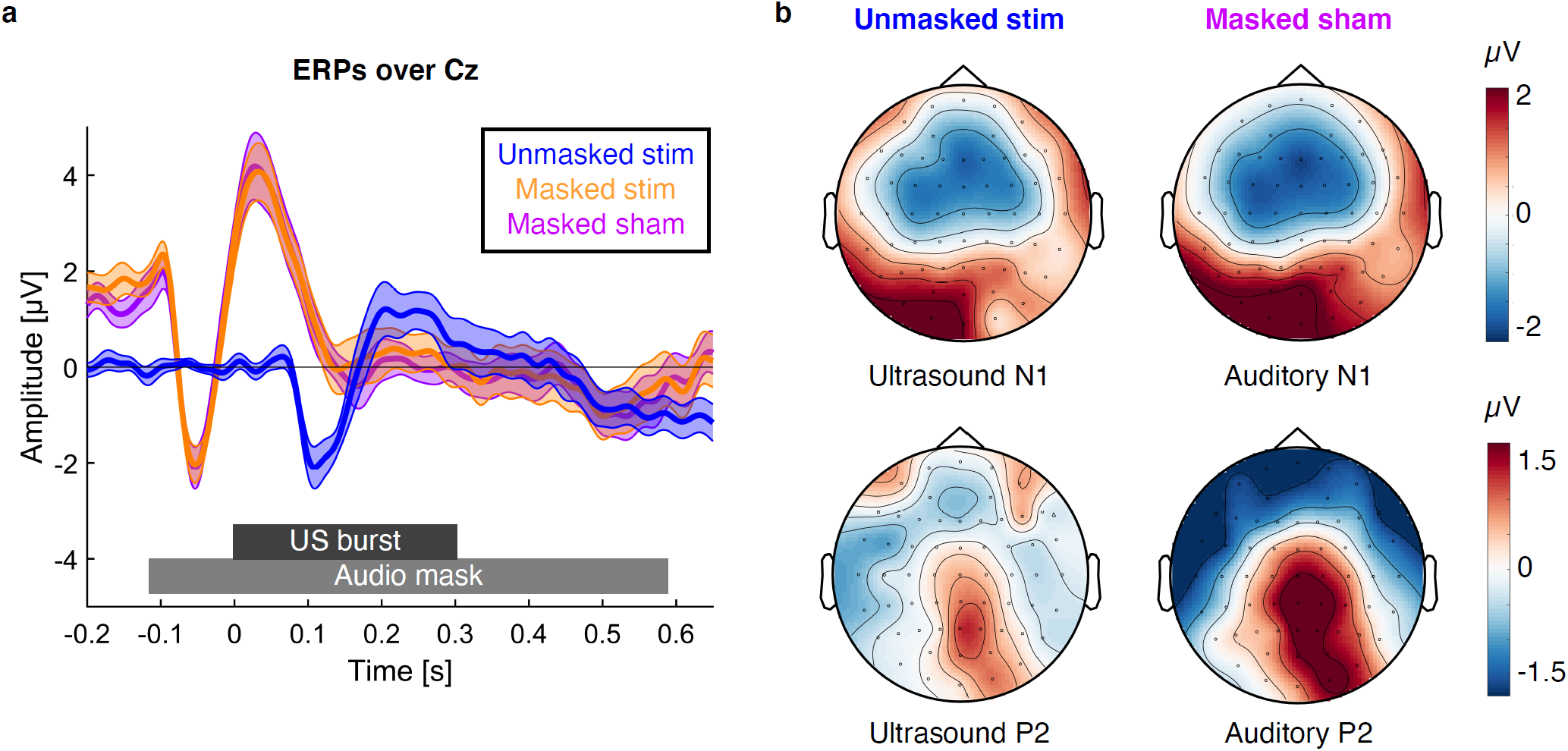
ERP results. a, ERP waveforms over Cz. ERPs over Cz for stimulation trials in the unmasked (blue) and masked (orange) condition as well as sham trials in the masked condition (purple) are shown. The unmasked ERP shows a response 100 ms after stimulation confirming that TUS induces an auditory ERP. The masked ERP shows a response to the masking audio tone starting 100 ms before the onset of the TUS burst. Shaded area represents standard error of the mean. **b, Comparison of Ultrasound ERPs and auditory ERPs**. US-N1 amplitude was defined as the mean amplitude 50 to 150ms after stimulation onset, whereas US-P2 amplitude was defined as the mean amplitude 150 to 250ms after stimulation onset (baseline window: −0.1 s to 0 s). Auditory N1 amplitude was defined as the mean amplitude −100 to 0ms before stimulation onset, whereas the auditory P2 component was extracted by taking the mean amplitude 0 to 100ms after stimulation onset (baseline window: −0.21 s to −0.11 s).

Ultrasound stimulation was delivered through a 500 kHz single-element geometrically-focused (64 mm aperture, 63.1 mm radius of curvature) transducer (H-107, Sonic Concepts Inc, Bothell, WA, USA). One stimulation trial consisted of a 300 ms burst applied at a 1 kHz pulse repetition frequency (PRF) and 50 % BDC as shown in Fig. 1a and 1b. The waveform was produced by an arbitrary waveform generator (Handyscope HS5, TiePie, WL Sneek, The Netherlands), amplified by a 55 dB broadband amplifier (1140LA, E&I, Rochester, NY, USA), and connected to the transducer via a matching network. The arbitrary waveform generator also generated a timing pulse which was recorded by the EEG to ensure synchronisation of ultrasound and EEG signals.

The ultrasound transducer was coupled to the scalp using a flexible polyurethane membrane (see Fig. 3a). This was fixed around the edges of the transducer using a custom mount with an o-ring seal. The cone was filled with degassed water through syringe connections on a back plate over the central circular transducer cut-out. Ultrasound coupling gel was liberally applied to the scalp of the subject before the coupling membrane was applied. The placement of the transducer was guided by a neuronavigation system (Brainsight; Rogue Resolutions; https://www.rogue-resolutions.com) with optical trackers placed on both the transducer and the subject.

**Fig. 3.**
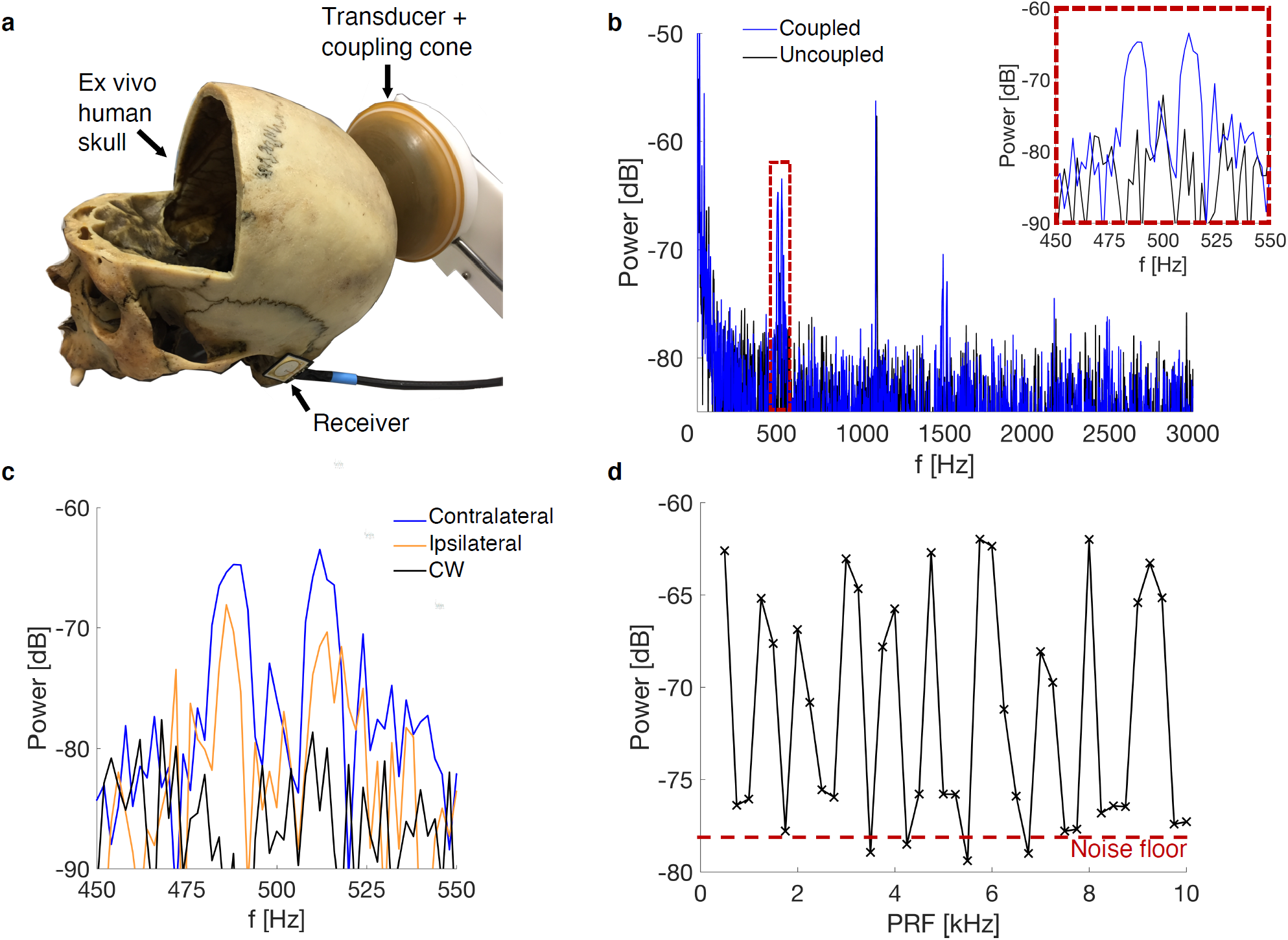
Vibration detection in an ex vivo human skull. **a**, Experimental setup: the TUS transducer was coupled posteriorly to an ex vivo human skull (the frontal skull portion had been removed for other purposes) via an expandable, water-filled coupling cone and US coupling gel. Vibrations were detected using a broadband receiver (700 Hz – 20 kHz) placed near the ear contralateral to the transducer. **b**, Power spectra for the received waveform with the receiver coupled and uncoupled to the skull. The applied US waveform was a 300 ms burst at a 50 % duty cycle and 500 Hz PRF. **c**, Recorded power spectra with the receiver positioned either contralateral or ipsilateral with respect to the transducer and for a continuous wave 300 ms US pulse. **d**, Power spectra amplitudes as a function of PRF. The maximum power in a 40 Hz window around the nominal PRF was extracted as the PRF was increased from 500 Hz to 10 kHz in steps of 250 Hz. The amplitude of the noise floor is indicated by the red dashed line.

#### EEG Recording and Preprocessing

EEG was recorded throughout the task from 54 Ag/AgCl sintered ring electrodes (BrainProducts 64-channel BrainAmp DC, BrainProducts GmbH, Gilching, Germany) arranged according to the 10%-system at a sampling rate of 500 Hz (low cutoff 10 s, high cutoff 250 Hz) referenced to FCz. EEG data were preprocessed and analysed using Fieldtrip [41] as well as in-house MATLAB scripts. EEG data were first epoched −1 s to 1.5 s around stimulation onset and visually inspected for artefacts. In order to identify ocular and muscle artefacts an independent components analysis was applied before the cleaned data were again visually inspected and trials containing remaining artefacts removed. Channels that had to be removed in order to allow room for the transducer to be coupled were subsequently reconstructed using a weighted neighbour approach as implemented in the fieldtrip toolbox [41] before re-referencing the data to common average reference, resulting in 60 electrodes in total.

#### Data Analysis

##### Behaviour

Behavioural data were analysed using in-house MATLAB scripts and JASP (JASP Team (2019). JASP(Version 0.9.2) [Computer software]). In order to assess participants’ ability to distinguish stimulation from sham trials, d’ values were calculated by subtracting normalised false alarm rates from normalised hit rates. A hit was defined as a stimulation trial that was correctly identified as such, whereas a sham trial that was falsely identified as an active stimulation trial was classified as a false alarm. Since d’ values for masked and unmasked trials deviated from normality (Shapiro-Wilk Test: unmasked: W=0.830, p=0.004; masked: W=0.603, p<0.001), one sample Wilcoxon signed-rank tests were conducted. Matched rank biserial correlations are reported as effect sizes for the Wilcoxon test. Masking success was further determined by conducting dependent samples t-tests comparing d’ values in the unmasked and masked condition. Effect sizes were estimated using Cohen’s d. Outliers were identified as values higher than Q3 + 1.5 times the interquartile range.

##### EEG

To investigate stimulation evoked effects, data were low-pass filtered at 30Hz and event related potentials (ERPs) were computed for every participant (baseline: −100 ms to 0 ms). In order to assess auditory activity associated with TUS, event-related potentials over EEG electrode Cz were extracted [42-44] and two main components analysed. US-P2 amplitude was defined as the mean amplitude 150-250 ms after stimulation onset while US-N1 amplitude as the mean amplitude 50-150 ms after stimulation onset. Mean US-P2 and US-N1 amplitudes were compared between conditions using repeated measures ANOVAs and follow-up dependent samples t-tests. Effect sizes for ANOVAs were estimated using partial eta squared (η^2^_p_), whereas Cohen’s d was used to estimated effect sizes for post-hoc t-tests.

### Skull audio recordings

Given the previously cited work in rodents [30,31], we hypothesised that auditory activation may be due to bone conduction: that is, as TUS passes through the skull it induces a radiation force on the skull at the modulation frequency which then propagates through the skull as a flexural wave to the cochlea, stimulating the standard acoustic pathway [45]. The possible coupling to flexural waves was investigated using an ex vivo human skull in air. In this instance, the ultrasound transducer was coupled to the skull with the coupling cone and US gel and a piezoelectric transducer (SPS-2220-03, Sonitron, Sint-Niklass, Belgium), with an active area 16 mm × 20 mm and bandwidth of 700 Hz to 20 kHz, was coupled to the skull close to the ear canal (see Fig 3a). All incident US bursts were 300 ms in length at a 50 % BDC and a PRF that was varied from 250 Hz to 10 kHz in steps of 250 Hz.

### Results

#### Behaviour

D’ results for unmasked and masked trials are shown in Figure 1c. Without auditory masking, participants were reliably able to detect stimulation trials (V=153, p<0.001, rank-biserial correlation=0.789). Masking the stimulation-induced sound with a masking tone resulted in detection rates not significantly different from zero (V=98, p=0.320, rank-biserial correlation=0.146). Masking success was further assessed by comparing d’ values between the two conditions directly. A dependent samples t-test revealed a significant reduction in d’ values when masking was applied (t(17)=4.352, p<0.001, Cohen’s d=1.026, 95% CI for the mean difference=[1.170 3.372]). Although the overall group d’ did not significantly differ from zero when masking was applied, three participants were still able to detect the stimulation with d’ values of 4.653, 3.804, and 3.407 (Fig. 1c). These three outliers were removed from all subsequent analyses. Only one participant of all 18 reported having seen phosphenes, and reported having seen these during both the unmasked and masked blocks. This participant was not one of the three outliers able reliably to detect stimulation during the masked block.

#### EEG

ERP waveforms over Cz are shown in figure 2a. In the unmasked condition, stimulation elicited a waveform characteristic of an auditory evoked potential (Fig. 2) [42-44] with a pronounced negative deflection around 100ms (US-N1: one-sample t-test: t(14)=-5.886,p=<0.001, Cohen’s d=-1.520, mean=-1.053, 95% CI=[-1.436 −0.669]; US-P2: one-sample t-test: t(14)=1.360, p=0.195, Cohen’s d=0.351, mean=0.670, 95% CI=[-0.386 1.726]). This response was abolished when auditory masking was applied. A 2×2 repeated-measures ANOVA with the factors MASKING (unmasked vs. masked) and COMPONENT (US-P2, US-N1) revealed a significant MASKING x COMPONENT interaction (F(1,14)=15.753, p=0.001, η^2^_p_=0.529). Post-hoc dependent-samples t-tests revealed that masking significantly affected US-N1 amplitude (t(14)=-5.279, p<0.001, Cohen’s d=-1.363, mean difference=-2.518, 95% CI=[−3.541 −1.495] while no difference in US-P2 amplitudes could be observed (t(14)=0.560, p=0.584, Cohen’s d=0.145, mean difference=0.423, 95% CI =[-1.196 2.041]).

In order to test whether any differences between stimulation and sham during masking can be observed, we conducted an additional 2×2 repeated-measures ANOVA with the factors STIMULATION CONDITION (stimulation vs. sham) and COMPONENT (US-P2, US-N1). This analysis only revealed a significant main effect of the factor COMPONENT (F(1,14)=9.816, p=0.007, η^2^_p_=0.412. No differences between stimulation and sham could be found (STIMULATION CONDITION x COMPONENT: F(1,14)=3.253, p=0.093, η^2^_p_=0.189).

#### Ex vivo skull recording

The experimental setup we used to investigate our hypothesis that auditory activation was due to bone conduction is shown in Fig. 3a. The signals recorded by the piezoelectric receiver were Fourier transformed and, for a 500 Hz modulation, peaks at 500 Hz were detected (Fig. 3b,c), consistent with the hypothesis that the 500 kHz signal is being absorbed and vibrating the skull at the modulation frequency. Peaks at the odd harmonics of the 500 Hz, i.e., 1500 Hz and 2500 Hz, were also detected which are consistent with what would be expected for a square wave modulation (Fig. 3b). When the receiver was uncoupled, the peaks were not present, indicating that the signal was not airborne. Interestingly, the spectra at each of the frequencies were split into two peaks approximately 10 Hz either side of the nominal frequency. The underlying explanation for this phenomenon is unknown.

Fig. 3c displays the recorded power spectra +/-50 Hz around the modulation frequency. To investigate whether peak amplitudes vary as a function of recording site, we compared amplitudes recorded from the contralateral ear canal to the power spectra of recordings at the ear canal ipsilateral to the transducer. This revealed amplitude variations of up to 6 dB, suggesting that skull topology may be important in the transmission of the flexural wave to the ear canals and may result in different sound localisations between participants. Moreover, the signal was not present for 300 ms continuous wave pulses confirming that the recorded waveform is a consequence of the modulating envelope (PRF). Fig. 3d shows the impact of the PRF on the amplitude of the received signals at each corresponding nominal frequency. For each burst, the PRF was increased by 250 Hz up to 10 kHz with the individual pulse lengths adjusted to maintain a BDC of 50 %. As in each case the spectrum was again split into two peaks, the maximum amplitude in a 40 Hz window around the driving PRF was extracted. This revealed variations in peak amplitude up to 12 dB, suggesting that the strength of the frequency response may be dependent on the modulation frequency as well as individual skull structure.

## Discussion

TUS is seen as the newest addition to a collection of methods that non-invasively stimulate the brain in order to investigate causal relationships between brain function and behaviour [1]. We investigated whether TUS can feasibly be applied to healthy human participants without the need for additional sham stimulation conditions. Eighteen healthy human participants received TUS to the visual cortex randomly intermixed with trials without stimulation. While only one subject reported having seen phosphenes during stimulation, we found reliable electrophysiological and behavioural evidence of auditory activation during TUS. ERP waveforms time-locked to stimulation onset resembled activity elicited by audible sound and all participants reported hearing an audible tone that enabled them to distinguish between stimulation and non-stimulation trials. Our findings indicate that sham stimulation conditions need to be adjusted to control for these unwanted auditory effects.

Auditory confounds during TUS have recently been reported in rodents with TUS eliciting nonspecific auditory activation that drives outcome while chemical deafening or transection of the auditory nerve abolished behavioural and cortical responses to TUS [30,31]. However, the exact mechanism behind these effects remains unclear. Gavrilov & Tsirulnikov [46] reviewed evidence from studies conducted in the 1970’s also indicating that human participants can perceive a sound during TUS that matches in pitch the frequency of the modulating envelope. In the present study, we also found evidence that modulation of the ultrasound signal may be responsible for the auditory confounds observed. In ex vivo skull recordings we could show that signals at the PRF and its harmonics can be received from the skull. These effects were not present when the receiver was removed from the skull. Our results suggest that it is coupling of the TUS into a physical wave, rather than direct neurostimulation, that results in audible sound.

Carefully constructing control conditions is vital in order to be able to draw causal conclusions about the role the manipulated brain process plays in behavioural outcomes. Controlling for nonspecific side effects of stimulation ensures that any effects observed can be attributed to the manipulation. Sham stimulation conditions have traditionally been tailored to the specific shortcomings of the technique used [34,35]. In TUS however, so far, most studies compared its effects on behaviour or neural processing to conditions in which stimulation was not applied [26-28], the transducer was tilted away from the head [24,25] or a disc with high acoustic impedance was placed between the transducer and the subject’s head [23]. These sham procedures would not induce the acoustic artefact reported here and are therefore not well suited to control for acoustic effects resulting from coupling through the skull. The present study, together with recent animal work [30,31], puts this practice into a new perspective. For TUS to be an effective tool to investigate the causal relationship between brain processes and behaviour, suitable sham stimulation conditions controlling for auditory effects have to be identified.

Our results show that simply not stimulating might not be the most optimal control condition and that future studies using this increasingly popular technique in humans need to take steps to control for auditory confounds. We blinded participants to the stimulation by delivering a masking sound at the PRF. While this was effective in most of our subjects and significantly reduced detection rates, three participants out of 18 could nonetheless tell the difference between active stimulation and sham. This may be the result of individual differences in resonance properties of the skull. Our ex vivo skull recordings revealed a strong frequency response during TUS which is likely skull (participant) and location specific as has been reported in the bone conduction literature [45]. Moreover, varying the PRF revealed that the amplitudes vary substantially at different frequencies. This variation is consistent with experiments on the frequency dependence of bone conduction on cadaveric heads [47]. We note that the use of an air-filled skull means the flexural wave speed will be different than for the case of the skull loaded by soft-tissue, which will alter the locations of the peaks. The result implies that adjustment of the PRF may change the amplitude of the induced wave, possibly dependent on the resonant modes of the skull structure [48]. Masking levels will therefore be subject specific. To minimise auditory activation, combined adjustment of masking level and PRF may therefore be necessary. This could be envisaged as a preliminary test before the start of the behavioural experiments, optimising the US waveform for each participant in order to reduce the impact of non-specific auditory activation.

Whether auditory masking is the most effective means to control for auditory confounds or whether control stimulation sites, or other sham conditions, would be more appropriate remains a task for future research. For quasi-continuous wave ultrasound waveforms, it has been shown that smoothing the leading and trailing edges of the pulse can be used to avoid auditory confounds in mice [32], and smoothed waveforms have been used successfully in macaques to modulate neuronal activity [49]. The approach in [49] employed a 12 ms rise time and fall time to smooth the transients of a quasi-continuous wave insonation, which is possible when the pulse has a duration much longer than the smoothing. However for the 1kHz PRF employed here, a 12 ms smoothing would attenuate the 0.5 ms long pulses and thus is not an effective approach to mitigate the acoustic confound. The masking employed here was narrowband, consistent with the critical band concept in auditory masking [50], where signals at different frequencies are not as effective at masking tones.

In conclusion, TUS has the potential to be a powerful, non-invasive brain stimulation technique, particularly given its advantages in terms of spatial accuracy and ability to target deep structures. However, auditory confounds should not be neglected and the results reported here provide further evidence for the need to optimise (sham) stimulation parameters.

## Abbreviations

BDC: burst duty cycle;
EEG: electroencephalography;
ERP: event related potential;
IQR: interquartile range;
PRF: pulse repetition frequency;
TUS: transcranial ultrasound stimulation

## Acknowledgments

This research was supported by the Medical Research Council [Clinical Scientist Fellowship MR/K010395/1 to C.R.B.)] and the John Fell fund [Grant number 163/11] awarded to C.R.B. J.B. was funded by an EPSRC studentship [EP/F500394/1].

